# CITE-Viz: Replicating the Interactive Flow Cytometry Workflow in CITE-Seq

**DOI:** 10.1101/2022.05.15.491411

**Authors:** Garth L. Kong, Thai T. Nguyen, Wesley K. Rosales, Anjali D. Panikar, John H. W. Cheney, Brittany M. Curtiss, Sarah A. Carratt, Theodore P. Braun, Julia E. Maxson

## Abstract

**Summary:** The rapid advancement of new genomic sequencing technology has enabled the development of multi-omic single-cell sequencing assays. These assays profile multiple modalities in the same cell and can often yield new insights not revealed with a single modality. For example, CITE-Seq (Cellular Indexing of Transcriptomes and Epitopes by Sequencing) simultaneously profiles the single-cell RNA transcriptome and the surface protein expression. The extra dimension of surface protein markers can be used to further identify cell clusters – an essential step for downstream analyses and interpretation. Additionally, multi-dimensional datasets like CITE-Seq require nuanced visualization methods to accurately assess the data. To facilitate cell cluster classification and visualization in CITE-Seq, we developed CITE-Viz. CITE-Viz is a single-cell visualization platform with a custom module that replicates the interactive flow-cytometry gating workflow. With CITE-Viz, users can investigate CITE-Seq specific quality control (QC) metrics, view multi-omic co-expression feature plots, and classify cell clusters by iteratively gating on the abundance of cell surface markers. CITE-Viz was developed to make multi-modal single-cell analysis accessible to a wide variety of biologists, with the aim to discover new insights into their data and to facilitate novel hypothesis generation.

**Availability and Implementation:** CITE-Viz installation and usage instructions can be found in the GitHub repository https://github.com/maxsonBraunLab/CITE-Viz

**Supplementary Information:** Down-sampled peripheral blood mononuclear dataset (Hao et al. 2021): https://bit.ly/3vxbhfW

## 1 Introduction

The development of methodology enabling the high-throughput profiling of transcriptomes at the single-cell resolution (scRNA-Seq) has revealed previously unappreciated levels of cellular heterogeneity (Macosko et al. 2015). Since then, newer scRNA-Seq assays have been developed to profile multiple macromolecules in the same cell. For example, CITE-Seq is a multi-omic variant of scRNA-Seq that captures the cell surface proteome using antibody-derived tags (ADT) (Stoeckius et al. 2017). Multi-omic assays like CITE-Seq add new dimensionality to the data and enable novel insights, but often require nuanced approaches to extract meaningful results from the data.

Data visualization and classification of cell clusters are commonly performed in single-cell analyses, but current methods fall short for multi-omic assays such as CITE-Seq. Single-cell visualization platforms like ShinyCell are often only compatible with one transcriptome assay (Ouyang et al. 2021), while SCHNAPPs emphasizes data preprocessing and lacks tools for cell cluster classification (Jagla et al. 2021). Cell cluster classification methods for CITE-Seq data include scGate (Andreatta et al.) and the Single-Cell Virtual Cytometer (Pont et al. 2020).

However, these programs lack data visualization tools and easy-to-understand quality control (QC) metrics that are essential to holistically evaluate a single-cell experiment (Figure 1A). We identified a need for a tool that can better visualize multi-omic single-cell data and classify cell clusters, while remaining accessible to biologists with minimal computational training.

**Figure 1.**
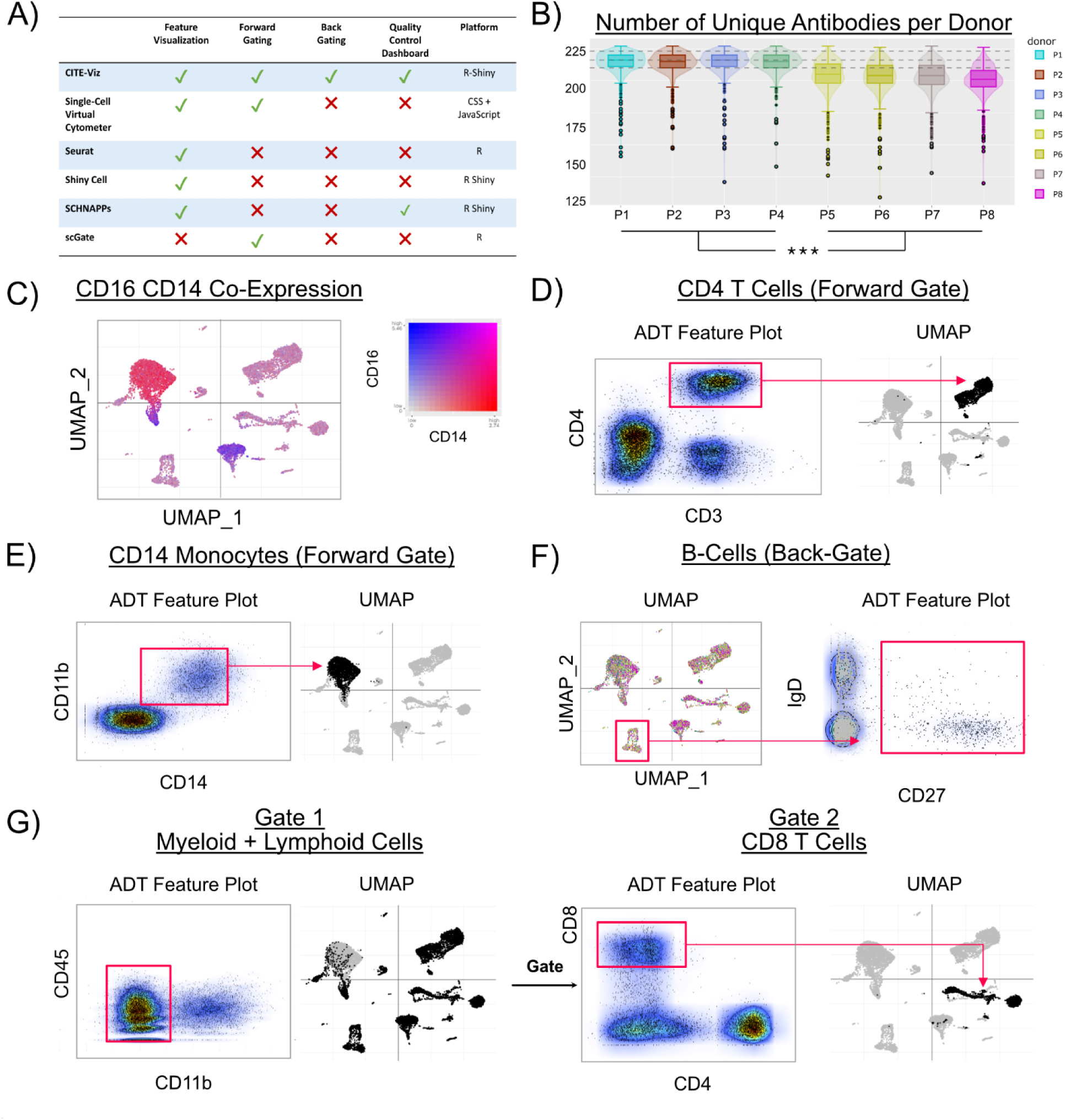
CITE-Viz suite of features. A) Comparison of CITE-Viz versus similar bioinformatics tools. B) One example of a QC metric, number of unique antibodies subset by patient donors, reveals lower unique counts in patients 5 – 8 compared to 1 – 4. C) Co-expression plot of ADT features CD14 and CD16 shows heterogeneous monocyte population. D) 1-layer gate that resolves CD4 T cells. E) 1-layer gate that resolves CD14 monocytes. F) Back-gate of B-cells shows IgD+ and CD27+/- cell population. Cells in the lower half of the red box represents naïve B-cells. G) 2-layer gate that resolves CD45- and CD11b- cells and then CD8 T cells.

Here we developed CITE-Viz, which enables users to visualize CITE-Seq data and classify cells by replicating the flow cytometry gating workflow. A flow cytometry workflow provides a familiar interface to bench biologists, and the nature of sequential gates can narrow specific cell populations of interest, which is difficult to do on a traditional UMAP. In real time, users can iteratively subset cell populations of interest using surface proteins, and see those cells reflected in the original dimension reduction (e.g. PCA, tSNE, UMAP) space (forward-gate). Conversely, cells can be selected in dimension reduction (DR) space and quickly located in a 2D gate window (back-gate). Additionally, CITE-Viz provides interactive quality control plots and 2D multi-omic feature expression plots. In conclusion, CITE-Viz is a single-cell, multi-omic visualization platform with a custom module that replicates the interactive flow cytometry gating workflow.

## 2 Methods

CITE-Viz was built on the R-Shiny platform to process Seurat-analyzed data. CITE-Viz accepts a preprocessed Seurat object in RDS format with one sample or an integrated object. After loading the data, QC metrics such as counts data per assay, unique ADT antibodies, and mitochondrial expression, can be subset by any categorical metadata in the user’s Seurat object (e.g. distribution of unique ADTs by patient donor sample).

The core of the flow cytometry workflow comes from a custom gate class. A gate class holds important metadata such as a custom gate label, X and Y axes labels (e.g. CD34, CD38, etc.), gate selection coordinates, input and output cell barcodes, etc. Most variables are intrinsic to a gate class except for the last, which is passed between gates. Gates are created by the simultaneous actions of a “Gate” button press and a rectangle selection of cells. Gates are saved using a list and can be downloaded to facilitate further analyses such as differential expression. Users can gate on any combinations of features and assays including SCT, RNA, ADT, etc.

## 3 Results

To demonstrate the utility of this program, CITE-Viz was used to re-analyze a peripheral blood mononuclear cell CITE-Seq dataset containing 211K cells and 225 antibody tags (Hao et al. 2021). For visualization purposes, the dataset was randomly down-sampled to 10K cells and then QC was performed. One example of a QC metric in CITE-Seq is the number of unique antibodies per sample. Using CITE-Viz, the unique antibodies per sample is clearly displayed and significant differences can be identified between patients 1 – 4 and patients 5 – 8 with a p_adj_ of 0 (Tukey HSD) (Figure 1B). To enable data exploration, CITE-Viz can also plot the relative expression levels of two features onto the UMAP simultaneously. An example is shown for CD16 and CD14 (ADT) markers, which reveals heterogeneous cell populations in the CD14 and CD16 monocytes (Figure 1C). This enables simultaneous exploration of two features in the dataset and enhances the data visualization capabilities of CITE-Viz.

Once markers of interest have been identified, cells can be classified by their expression of these markers using one or more “gates” in a flow cytometry-like workflow. An example of the classification of cell clusters with one gate is shown in Figure 1D, where cells with CD3+ and CD4+ surface protein signature represent a CD4 T cell population. Likewise, the same monocyte population from Figure 1C can also be orthogonally defined via the gating module using the CD11b+ and CD14+ markers (Figure 1E). Another type of one-layer gate is the back-gate unique to CITE-Viz, where selected cells in DR space are highlighted in the ADT space. For example, a selection of B-cells from the UMAP space shows a distinct set of cells with an IgD+ and CD27+/- signature (Figure 1F). To demonstrate serial gating, an initial CD45- and CD11b-gate selected a mixture of myeloid and lymphoid cells. From this gated cell subset, CD8+ and CD4-cells were identified to as CD8 T cells, which are visualized as a distinct cell cluster on the UMAP (Figure 1G).

## 4 Discussion

CITE-Viz is a single-cell multi-omic data visualization platform with an interactive module to help classify cell clusters. Users can see QC metrics clearly displayed and view multi-omic feature expressions with a 2-dimensional color matrix. Similar to flow cytometry, users can gate cells with antibodies (forward-gate) or gate from DR space (back-gate) in real time with minimal computational proficiency. The application of CITE-Viz to a peripheral blood mononuclear cell CITE-Seq dataset provided an orthogonal approach that was congruent with previously identified cell cluster identities (Hao et al. 2021).

A limitation of CITE-Viz pertains to data sparsity where gating by gene expression results in a high rate of data dropout. However, we speculate the development of better multi-omic sequencing assays will help resolve heterogeneous cell populations. Therefore, CITE-Viz was designed to gate cells based on any combination of assays and features to accommodate impending changes in sequencing technology. In conclusion, CITE-Viz provides a new user-friendly tool to visualize data and classify cell clusters in CITE-Seq data by replicating the iterative gating workflow of flow cytometry.

## Supporting information

Per

## Data availability

### Data Availability Statement

The data underlying this article are available in GEO (Gene Expression Omnibus) at https://www.ncbi.nlm.nih.gov/geo/, and can be accessed with GSE164378. The down-sampled data via the supplementary material section.

## Acknowledgements

We thank the following Oregon Health and Science University core facilities for their assistance: ExaCloud Cluster Computational Resource and the Advanced Computing Center.

## Funding

This work was supported by the American Society of Hematology Research Restart Award; American Society of Hematology Scholar Award; National Cancer Institute [K08 CA245224]. Funding for SAC was provided by the American Society of Hematology Research Restart Award; Collin’s Medical Trust Award; Medical Research Foundation Early Clinical Award; and the National Cancer Institute [F32CA239422].

## Conflicts of Interest

none declared.

## Notes

https://github.com/maxsonBraunLab/

### Competing Interest Statement

The authors have declared no competing interest.

